# Improved basic cytogenetics challenges holocentricity of butterfly chromosomes

**DOI:** 10.1101/2022.03.11.484012

**Authors:** Bernard Dutrillaux, Anne-Marie Dutrillaux, Mélanie McClure, Marc Gèze, Marianne Elias, Bertrand Bed’hom

## Abstract

Mitotic chromosomes of butterflies, which look like dots or short filaments in most published data, are generally considered to lack localised centromeres and thus to be holokinetic. This particularity, observed in a number of other invertebrates, is associated with meiotic particularities known as “inverted meiosis”, in which the first division is equational, i.e., centromere splitting-up and segregation of sister chromatids instead of that of homologous chromosomes. However, the accurate analysis of butterfly chromosomes is difficult. 1) Their size is very small, equivalent to a single band of a mammalian metaphase chromosome. 2) They lack satellite DNA/heterochromatin in putative centromere regions and therefore marked primary constrictions. Our improved conditions for chromosome preparations in six butterfly species belonging to the Nymphalidae and Pieridae families challenges the holocentricity of their chromosomes: in spite of the absence of primary constriction, sister chromatids are recurrently held together at definite positions during mitotic metaphase, which makes possible to establish karyotypes composed of acrocentric and sub-metacentric chromosomes. The total number of chromosomes per karyotype is roughly inversely proportional to that of non-acrocentric chromosomes, which suggests the occurrence of frequent Robertsonian-like fusions or fissions during evolution. Furthermore, the behaviour and morphological changes of chromosomes along the various phases of meiosis do not differ much from those of canonical meiosis. In particular at metaphase II, chromosomes clearly have two sister chromatids, which refutes that anaphase I was equational. Thus, we propose an alternative mechanism to holocentricity for explaining the large variations in chromosome numbers in butterflies: 1) in the ancestral karyotype, composed of about 60-62 acrocentric chromosomes, the centromeres, devoid of centromeric heterochromatin/satellite DNA, were located at contact with telomeric heterochromatin; 2) the instability of telomeric heterochromatin largely contributed to drive the multiple chromosome rearrangements, which occurred during butterfly evolution.

## Introduction

Lepidopterane of the largest insect orders. Traditionally, they have been divided in 2 groups, based on morphological and ecological features: butterflies and moths. While butterflies form the superfamily Papilionoidae, which comprises about 18,000 species, moths do not form a monophyletic group of species. Nevertheless, the distinction between butterflies and moths remains practical. The haploid genome of Lepidoptera, with a mean C-value of 0.66 ± 0.04 pg of DNA, is one of the smaller amongst insects, and amongst Lepidoptera, the mean size of butterfly genome (0.4 pg) is almost half that of moth genome (0.7 pg) [calculated from Gregory, 2003, 2011]. This suggests that some differences exist in the chromosome structure and composition of these 2 groups, but the question remains open because cyto-molecular studies were essentially limited to moths, which have a larger economic importance than butterflies [Wolf et al., 1997, Mediouni et al., 2004, Fukova et al., 2005]. Compared to Mammalian genome (± 3.5pg), the small size of the Lepidoptera genome is partly due to the relative paucity in the various types of repeated DNA sequences, including transposable elements [d’Alençon et al., 2010]. In spite of their small genome, their modal number of chromosomes (2N = 62) is among the highest in animals. A direct consequence is that butterfly chromosomes are very small, about 1/15 ^th^ of medium sized Mammalian chromosomes, and lack structures harbouring DNA repeats such as primary constrictions (centromeric heterochromatin) and euchromatin banding. Most cytogenetic data on butterflies were performed on male gonads and the techniques often privileged squashes, or direct fixation without previous hypotonic shock and colchicine treatment. In such conditions, mitotic chromosomes often look like dots at metaphase and short shapeless filaments at pro-metaphase. The difficulty to observe distinct sister chromatids and primary constriction led to the notion that the chromosomes of butterflies lack localized centromeres and are thus holokinetic, a particularity reported in other insect orders such as Dermaptera, Ephemeroptera, Hemiptera, Heteroptera, Homoptera, Odonata, Psocoptera, Thysanoptera and Trichoptera [Mola and Papeschi, 2006, for review]. Comparisons of DNA sequences provide convincing evidence that these orders are scattered across the phylogenetic tree of insects [Kjer et al., 2016, Misof et al., 2014, Drinnenberg et al., 2014]. Thus, holocentricity has not evolved once, but is rather derived after multiple convergent events from the widely spread centric chromosomes [Bauer, 1967, Melters et al., 2012]. In butterflies as in other taxa, the holocentric nature of chromosomes was confirmed by the lack of centromeric histone H3 variant CenH3 [Drinnenberg et al., 2014]. In spite of real advances, as in the nematode *Caenorhabditis elegans* [Howe et al., 2001, Zedec and Bures, 2012], the structure of holocentric chromosomes remains largely unknown, in particular in insects, and there is no strong argument to consider that identical structures replaced localised centromeres in the various taxonomic groups which independently acquired holocentricity during evolution. Consequently, holochromosomes may differ from order to order, and even among various taxa inside a given order, as butterflies and moths amongst Lepidoptera. Another difficulty is that the appearance of chromosomes may depend on several parameters such as their size, the phase in the cell cycle, the tissue studied and the technique used. It is well known that the primary constrictions marking the centromeres, hardly visible at early pro-metaphase, becomes obvious at metaphase, when chromatids are condensed and cohesins are cleaved. Similarly to butterflies, other insect cytogenetic studies were generally performed on male germinal cells [Smith and Virkky, 1978]. Working on both beetles and mouse male germinal cells, we were surprized to occasionally observe similar atypical morphologies of their monocentric chromosomes. A more systematic study was then performed on mouse spermatogenesis, which showed that the appearance of mitotic chromosomes deeply changes along divisions from gonocytes to late spermatogonia [Coffigny et al., 1999]. While chromosomes have hyper-cohesive, long and thin chromatids with hyper-methylated DNA in gonocytes, they become shorter with more fuzzy and separated chromatids concomitantly with their hyper-methylation loss in spermatogonia [Bernardino et al., 2002]. The strong relationship between DNA methylation status and chromatid compaction and cohesion remained unexplained, but the similarity of transient morphological changes of chromosomes in both mouse and beetles (personal data), suggests that they may occur in a large range of animals. These tissue and developmental stage changes may render difficult the interpretation of chromosome morphology in germ cells. Electron microscopy also showed that the size of the kinetochore plate could vary at different stages of gametogenesis [Wolf et al., 1997].

During meiosis, the behaviour of monocentric and holokinetic chromosomes was found to be different. Meiosis is described as inverted in species with holochromosomes [Mola and papeschi, 2006, Lukhtanov et al., 2018]: at metaphase I, bivalents are disposed perpendicularly to the spindle, each homolog is attached to several spindle fibres and sister chromatids segregate at anaphase I (pre-reduction) instead of homologous chromosomes as in the post-reduction of canonical meiosis in species with monocentric chromosomes. Consequently, the typical morphology of chromosomes at metaphase II (MII), with 1 centromere linking 2 well-separated fuzzy sister chromatids should not be observed in species with true holochromosomes, but the difficulty to observe the morphology of the small MII butterfly chromosomes prevented to assess the occurrence of inverted meiosis. The presence of holochromosomes was questioned in some species and monocentric mitotic chromosomes with “G-banded” chromatids were even described in Pieridae [Bigger, 1975, Rishi and Rishi, 1977, 1979], but the photos shown were much less convincing than the drawings, a flaw also shared by many reports on holochromosomes. With the purpose of studying the karyotypes of some butterfly species, we adapted our usual technique to this particular material [McClure et al., 2018]. After some additional improvements, we describe here a very simple technique for the study of Lepidoptera chromosomes, particularly adapted to small cell samples. It allowed us to observe not only atypical monocentric chromosomes in mitotic cells, but also figures hardly compatible with the occurrence of an inverted meiosis. Chromosomes of some selected species of butterflies are shown and compared to those of *Drosophila melanogaster*, whose chromosomes are well known and DNA content not very different. The meaning of these basic cytogenetic observations is discussed.

## Material and methods

### Species

*Drosophila melanogaster* Melgen, 1930, obtained from a laboratory strain was selected for chromosome size comparisons.

Butterfly species included in this study are the following:

*Anaea eurypyle* Hubner, 1819 (Pointed leafwing, Nymphalidae, Charaxinae) is a Neotropical species. Specimens were obtained as pupae from a butterfly farm. This species was selected amongst those we studied because its chromosome number of 62 is presumed to be that or very close to that of butterfly ancestors.

*Pieris brassicae* Linnaeus, 1758 (Large White, Pieridae) was caught at Bois-le-Roi, France (48.28.25 N, 02.41.50 E). This species was selected because the holocentry of its chromosomes was previously questioned [Bigger, 1975, Rishi and Rishi, 1977].

*Pieris rapae* Linnaeus, 1758 (Small White, Pieridae), caught at Bois-le-Roi, was selected because its genome size is known.

*Archaeopreponia demophon* Linnaeus, 1758 (One-spotted Demophon, Nymphalidae, Charaxinae). This neotropical species, obtained as pupae from a butterfly farm, was selected for its small number of chromosomes.

*Melinaea menophilus* n. ssp Hewitson, 1856 (Hewitson’s tiger, Nymphalidae, Danainae). Specimens of this neotropical species, obtained from eggs laid in captivity by wild-caught females, were also selected for their small chromosome number.

*Ithomia salapia aquinia* Hopffer, 1874 (Nymphalidae, Danainae). This neotropical species was selected among those studied because its chromosomes exhibit many C-bands. Specimens were obtained from eggs laid in captivity by wild-caught females.

### Methods

Dividing cells were obtained from either cerebral ganglia of caterpillar (*Pieris* species) or testes from freshly killed male imagine (all species). All dissections were performed on a glass slide in 1 or 2 drops of a solution of 0.88 g KCl in 100 ml distilled water. **Important remark:** fresh KCl powder is not stable and becomes more or less quickly hydrated, especially in the field. We deliberately used hydrated KCl, more stable for weighing. All centrifugations were performed at about 7 g. for 7 min. After all centrifugations and settling, cells were suspended by gently taping the tube. Carnoy I fixative was used for all fixations. Staining was performed in 2% Giemsa in tap water for 7-10 min, and occasionally with DAPI. C-banding was performed as described by Angus [1982].

#### Cerebral ganglia and eggs

immediately after dissection (ganglia cells), they were placed and ruptured inside an Eppendorf tube where they were maintained in 1 ml of the KCL 0.88 g/L solution added with colchicine (5 μL of a 4 mg/L solution) for 45 min. After centrifugation, the supernatant was replaced by either an aqueous solution of KCl 0.55g/L or foetal calf serum diluted in water (1vol: :4 vol) for 10 min. One drop of Carnoy fixative was added just before centrifugation. The pellet obtained was not always visible and about 100 μL was left in the tube before immediate addition of 1 mL fixative. After a new centrifugation, about 75-100 μL of supernatant were left in which cells were gently suspended with a Pasteur pipette, and dropped from about 20 cm on glass slides special for FISH.

#### Testicle

Immediately after dissection, the unique testicle was placed in an Eppendorf tube containing 0.6 mL of an aqueous solution of KCl (0.85 g/L), in which colchicine was eventually added, as above. The testicle was then ruptured using a piston adapted to the internal diameter of the tube, which was gently turned during about 10 seconds. For salvaging adhering cells, the piston was rinsed with 2-3 drops of the 0.85 g/L KCl solution, of which 0.5 mL was added. The tube was gently taped, let for 1 h, taped again and one drop of fixative was added. Centrifugation, fixation and spreading of the cells were as above.

Confocal microscopy: We employed a recent method developed by Zeiss corporation for studying specimen in thin sections, using confocal scanning laser microscopy and image processing software that allows us to produce high resolution three-dimensional visualizations. We acquired confocal images with a Zeiss LSM 880 laser-scanning confocal microscope using a 63 X/1.4 oil DIC objective. The DAPI fluorescence signal was collected with an Airyscan head using a 32 GASP detector array (Quantum efficiency (QE) of the detector was about 50%) in super resolution mode, which increases the SNR and the resolution by a factor 1.7.

DAPI fluorescence of the samples was excited by the 405 nm diode laser (power 1% with pixel dwell 2.52 µs). Emission was collected with a band pass filter 420-480 + 495-550 nm. Images were acquired with 16-bit depth and 0.2 airy unit for each elementary detector of the Airyscan head and recorded with pixel size of 35 nm. Typically producing 11 slice z-stacks comprising of individual focal planes, each separated by a 160 nm z step, corresponding to a z depth 1,75 μm. The fluorescence signal from each z-plane was projected onto a maximum projection image by the software Zen Black version 2.3 (Zeiss corporation). Images were processed using the Fiji software [Schindelin et al., 2012].

## Results

### *Drosophila melanogaster* Melgen, 1830

Its well-known male diploid karyotype (2N = 8), obtained with a technique close to that of butterflies, is shown for comparison in Fig. 1a. Its diploid DNA content is 279 Mb (Mega base pairs) [Adams et al., 2000] (Table 1). C-banded heterochromatin, which represents about 1/6^th^ (chromosome 2) to 9/10^th^ (chromosomes 4 and Y) of whole chromosome lengths, largely corresponds to regions of sister chromatid joining, independently of the immediate centromere proximity (Fig. 1b). At metaphase/anaphase transition (Fig. 1c), heterochromatin disjoins but sister chromatids remain loosely attached, presumably by their centromeric regions.

**Table 1.** Comparison of diploid genome in Million base pairs (Mbp) and medium sized chromosome DNA content of 4 reference species: a medium sized chromosome of *P. rapae* is about 15 fold smaller than a human chromosome 8 (HSA 8); 4 fold smaller than a *D. melanogaster* chromosome 2 (DME 2) and 3 fold smaller than an average chromosome of *B. mori* a:[Shen et al, 2016], b:[Carvalho, 2002].

| Species | Whole genome |  | Average chromosome (HSA8, DME2) |  |  |
| --- | --- | --- | --- | --- | --- |
|  | Diploid chromosome Number | Diploid DNA content (Mbp) | Size (Mbp) | % HSA | % DME |
| *Homo sapiens* | 46 | 6400 | 145 (8) | 100 | 330 |
| *Bombyx mori* | 28 | 864 (a) | 31 | 21 | 70 |
| *Pieris rapae* | 50 | 492 (a) | 10 | 7 | 23 |
| *Drosophila melanogaster* | 8 | 279 (b) | 44 (2) | 30 | 100 |

**Fig. 1.**
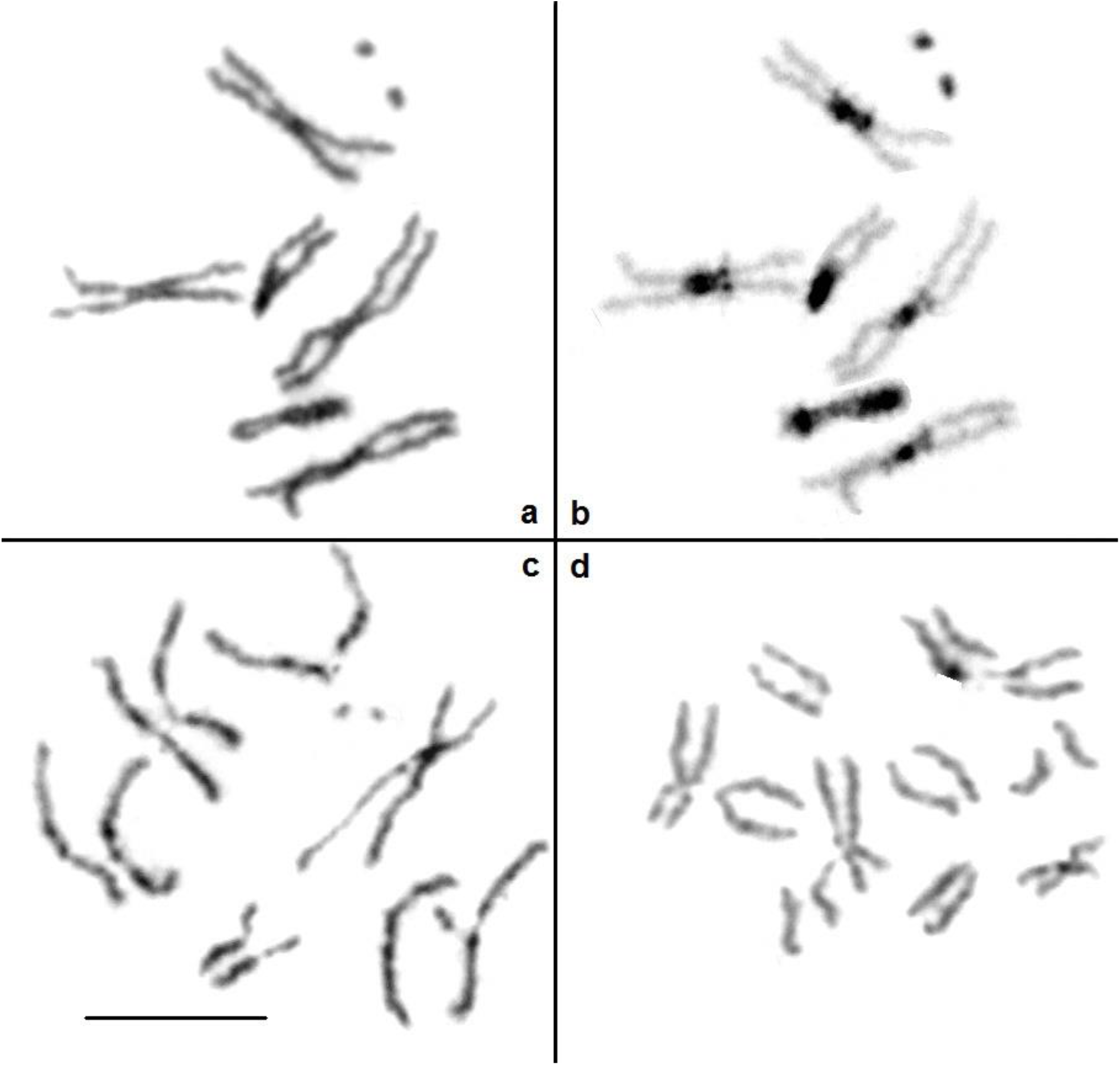
(a, b, c) *Drosophila melanogaster* mitotic chromosomes: same pro-metaphase after Giemsa staining (a) and C-banding (b) and early anaphase (c). (d) Part of a Giemsa stained early anaphase of a *Pieris brassicae* caterpillar cerebral ganglia cell, in which indisputable primary constrictions exist on both acrocentric and sub-metacentric chromosomes. Bar = 10 μm for a, b, and all other figures but 5 μm for 1d.

### Anaea eurypyle

The male karyotype is composed of 62 chromosomes (62,ZZ), the very small size of which makes difficult the analysis of their morphology. However, most of them look acrocentric (Fig. 2), alongside with some sub-metacentric chromosomes. At diakinesis, all but 2 bivalents display 1 chiasma, most frequently in interstitial position.

**Fig. 2.**
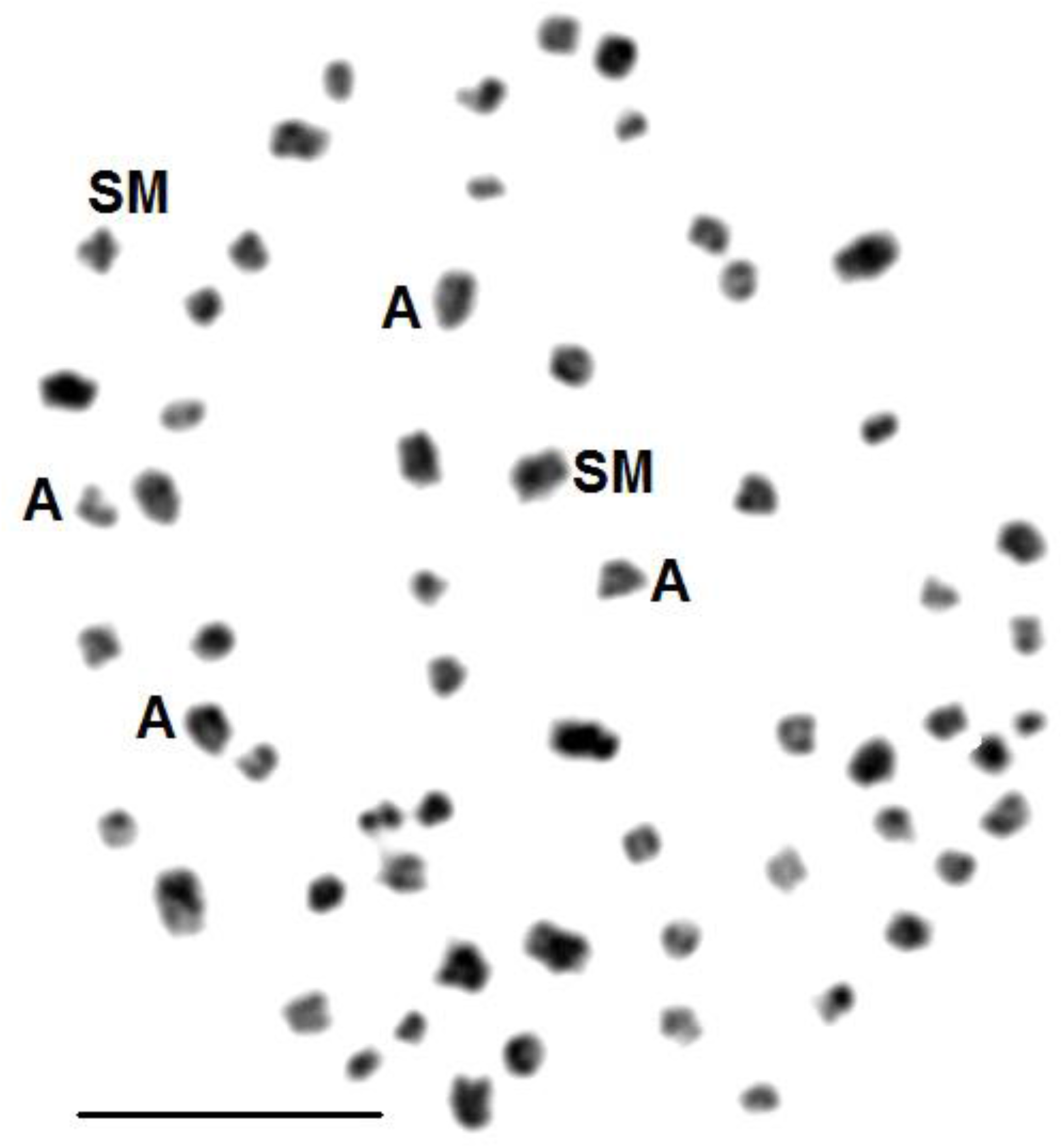
Mitotic metaphase of *Anaea eurypyle*. Most chromosomes look acrocentric and a few look sub-metacentric (some of them are labelled A and SM, respectively). The DNA content of each *D. melanogaster* metacentric autosome (Fig. 1) is equivalent to that of about 10 *A. eurypyle* chromosomes.

### Pieris brassicae

As previously described [Bigger, 1975] the male karyotype is composed of 30 chromosomes: 30,ZZ. The discrimination of sister chromatids in mitotic chromosomes is easier in caterpillar ganglia cells and eggs, but is occasionally possible in germ cells. After Giemsa staining, chromosomes generally have no clear primary constriction, i.e., no lighter staining regions but they often display 1 region where sister chromatids are brought closer together, which suggests the presence of localized centromeres (Fig. 3 a, b and Fig. I and II in complementary data). Somatic and germ cells roughly display similar chromosome patterns, with 7 sub-metacentric and 8 acrocentric chromosomes (Fig.3 a,b,c). The presence of monocentric chromosomes is clearer at metaphase/anaphase transition: at this stage, the morphology of *D. melanogaster* and *P. brassicae* chromosomes is quite similar (compare Fig. 1c and 1d). More or less discreet C-bands on either one or both homologues at many terminal but no interstitial regions indicate the presence of polymorphic juxta-telomeric heterochromatin. In some eggs or ganglia cells, 1 chromosome is almost entirely C-banded (Fig. III, complementary data). This particularity is not seen in homogametic ZZ spermatogonia, which may indicate that the W chromosome of ZW females is largely composed of C-banded heterochromatin, which fits with the richness in DNA repetitive elements of the W described in some moths and butterflies [Sahara et al., 2012]. In meiotic cells at diakinesis /metaphase I, a majority of bivalents (10-12/ 15) have a ring configuration (2 chiasmata). All have at least 1 terminal chiasma, and about half of the second chiasmata are interstitial. At metaphase I/anaphase I transition, chromosomes are very small and remained attached by the extremity of 1 or 2 chromatids but they have not the side-by-side position described for holochromosomes (complementary data Fig. IV). At metaphase II, chromosomes remain very small and their compaction is unusual. Their sister chromatids are distally well separated, fuzzy, and lightly stained, but clearly linked at a position of locally highly compacted chromatin. They form sub-metacentric and acrocentric chromosomes in numbers comparable to those of mitotic divisions (Fig. 3c). To further explore chromosome morphologies, DAPI stained mitotic cells at metaphase/anaphase transition were analysed through a high-resolution confocal microscope, which evidenced discreet chromatin amounts linking the sister chromatids at the place of their putative centromere (Fig. 4).

**Fig. 3.**
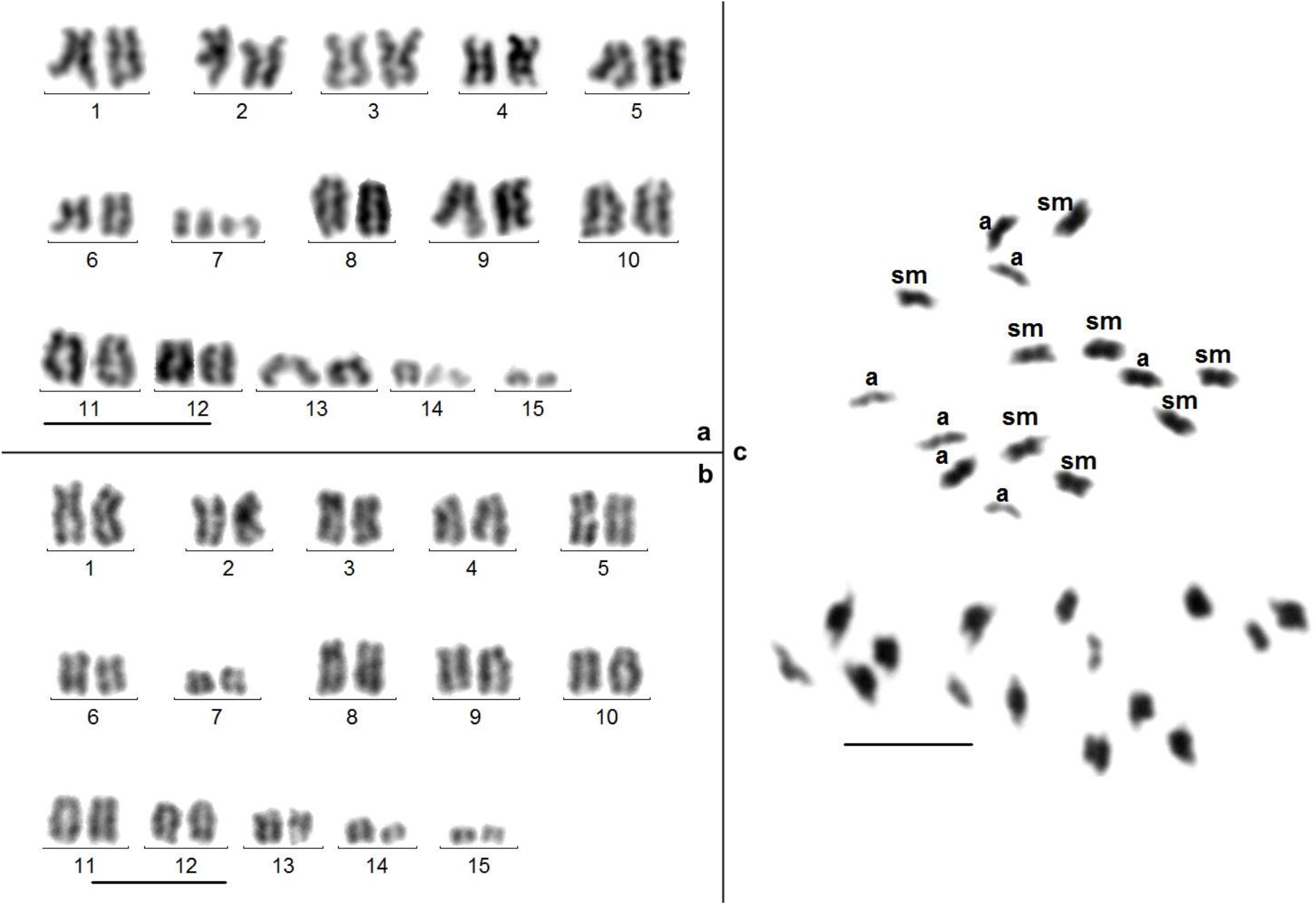
*Pieris brassicae*. a and b) Karyotypes from caterpillar brain cells. Sex chromosomes are not identified. c) “Brother” spermatocytes II displaying acrocentric (a) and sub-metacentric (sm) chromosomes. Their morphology, with strongly stained and highly compacted proximal and poorly stained fuzzy distal regions, seems to be typical for butterflies.

**Fig 4.**
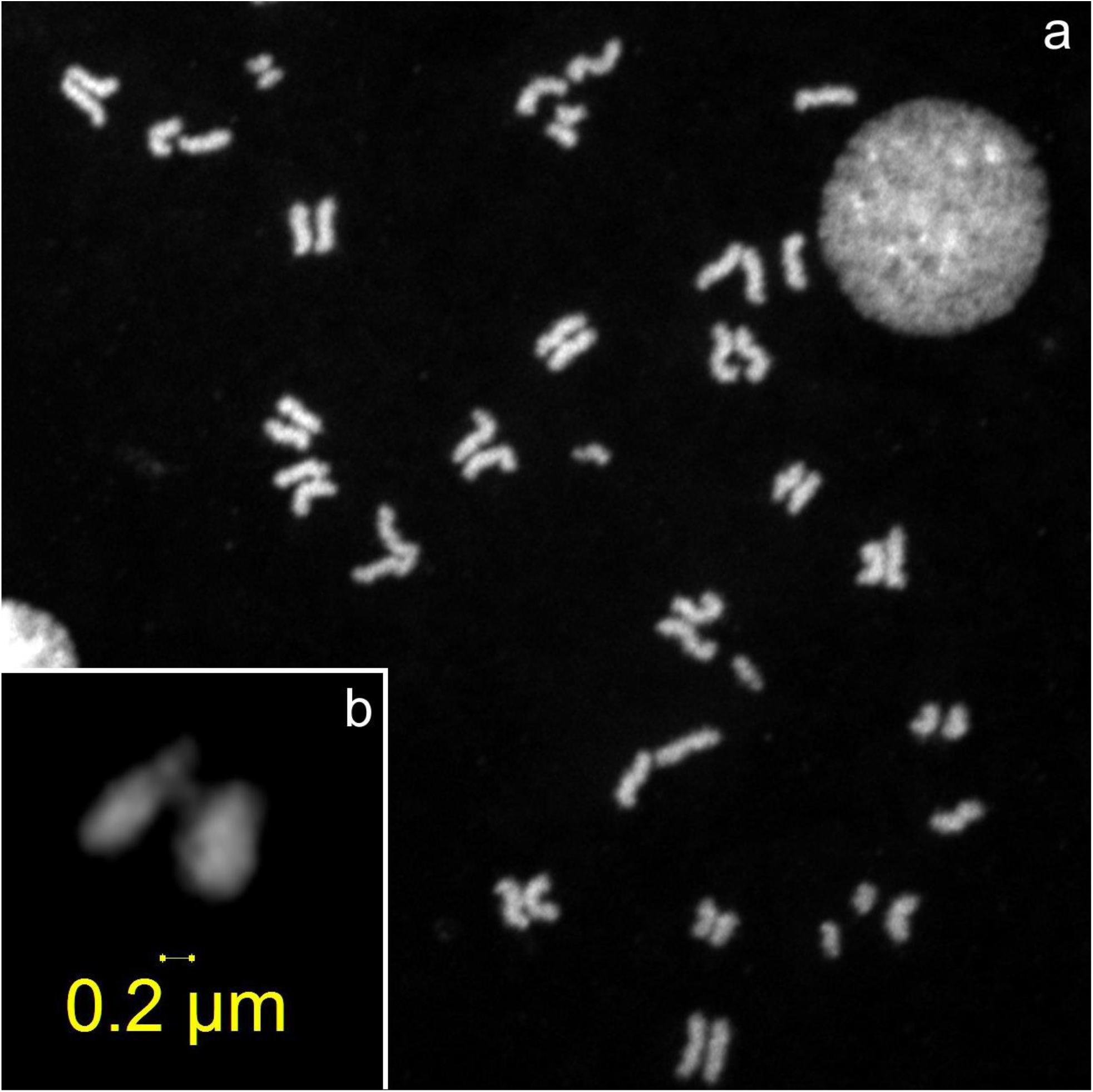
a) Confocal image of a DAPI stained ganglion cell at metaphase-anaphase transition. b) higher magnification of a small acrocentric chromosome from another cell.

### Pieris rapae

The male karyotype is composed of 50 chromosomes: 50,ZZ. Although the small size of chromosomes makes the analysis difficult, most mitotic chromosomes look acrocentric, and thus monocentric (Fig. 5a). The acrocentric morphology of many chromosomes is confirmed at metaphase I/anaphase I transition, when the homologues begin splitting (Fig. 5b). This stage contradicts the occurrence of an inverted meiosis.

**Fig. 5.**
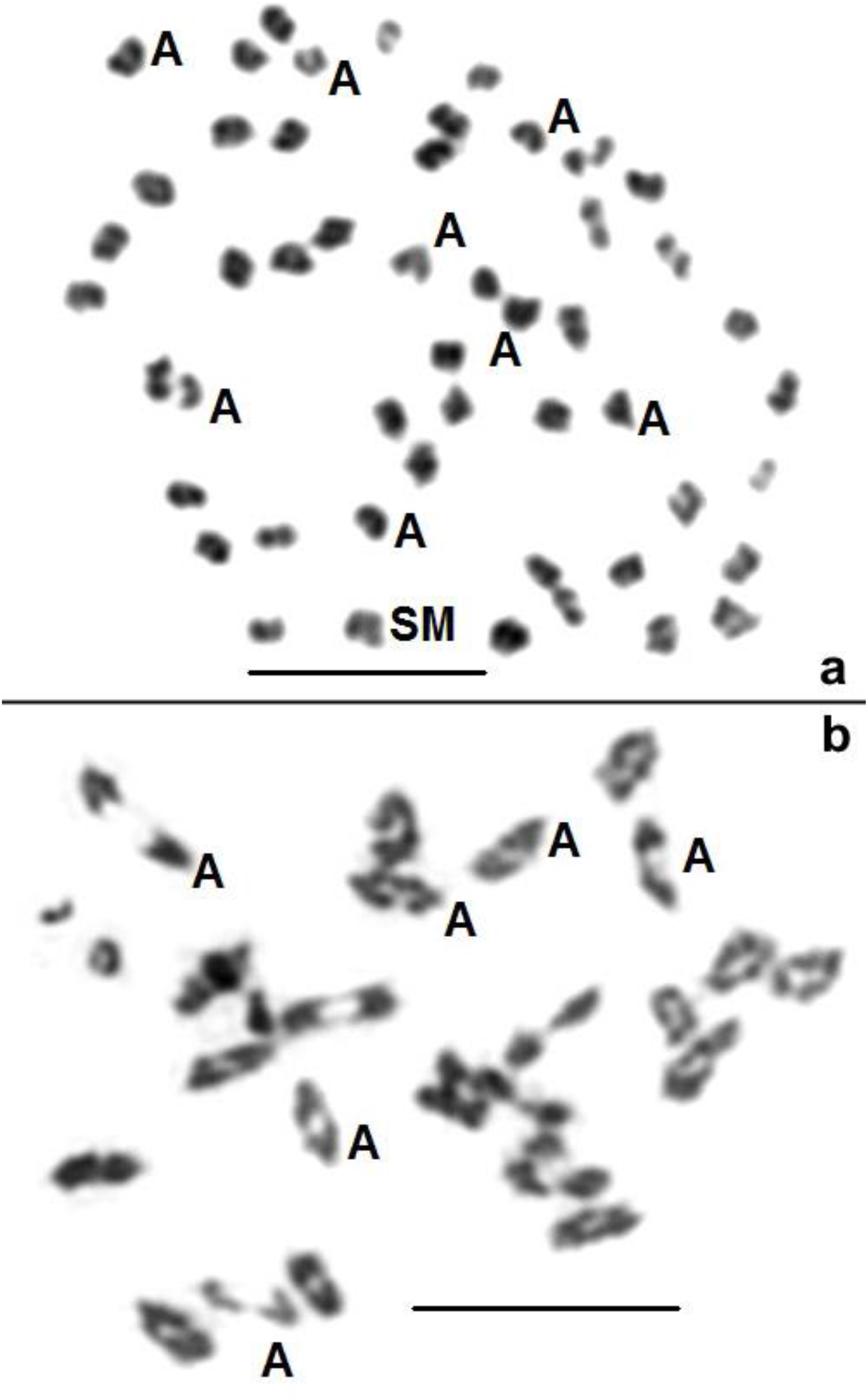
*Pieris rapae*. a) Metaphase of a caterpillar brain cell. Most chromosomes look acrocentric, some are indicated (A). b) Metaphase I/anaphase I transition. Homologs in most bivalents are in end-to-end association. Some of them are more or less dissociated and look acrocentric and pulled by a single point, their centromere, and not by multiple points along the chromatids, as expected for holochromosomes.

### Archaeoprepona demophon

The male karyotype is composed of 32 chromosomes: 32,ZZ, including 11 pairs of sub-metacentrics (Fig. 6a). At diakinesis/metaphase I, most bivalents display 1 interstitial chiasma (not shown). At metaphase II, largely separated sister chromatids are pale and fuzzy, whereas they look compacted at their junction, which makes a pseudo C-banding (Fig. 6b).

**Fig. 6.**
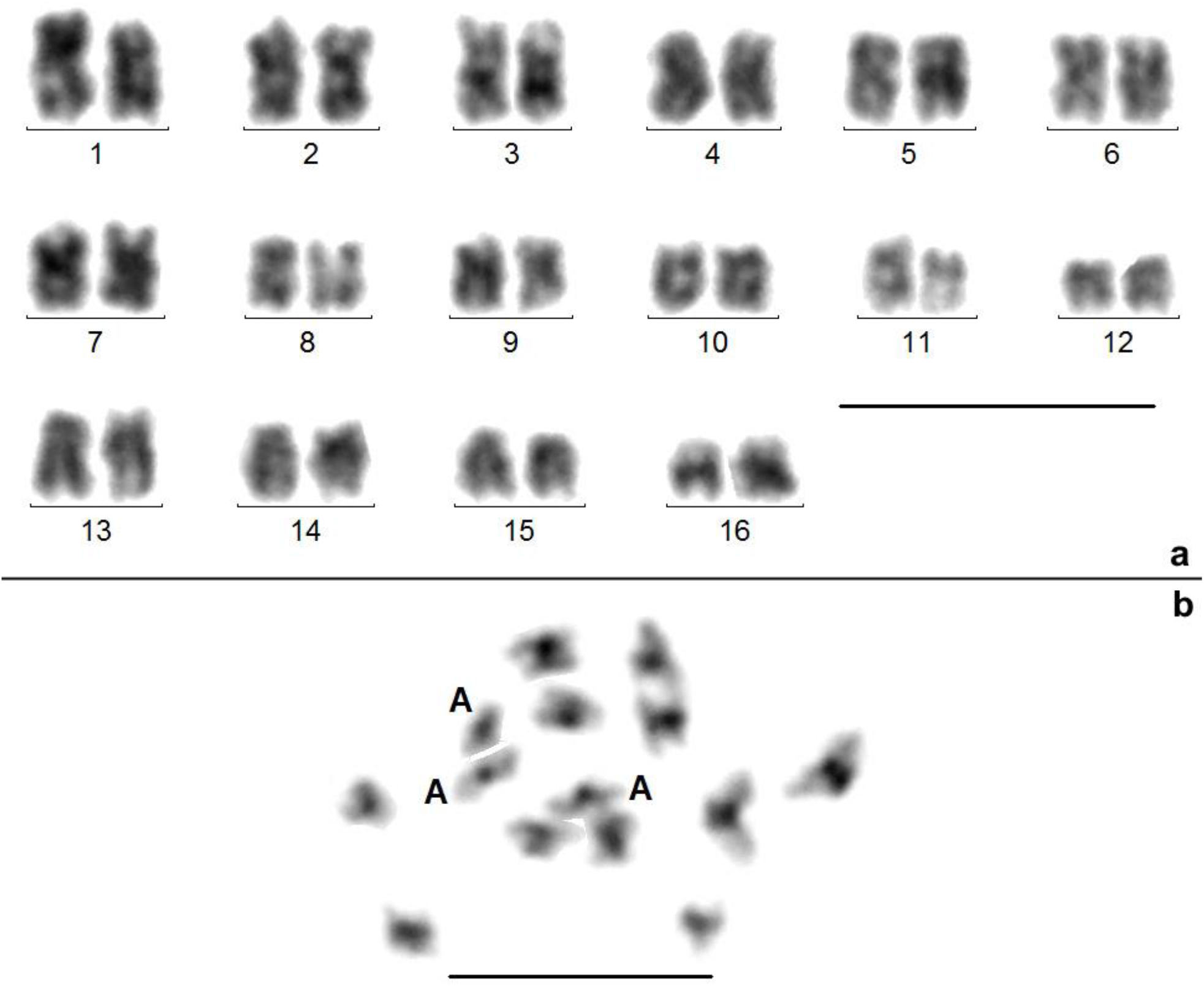
*Archaepreponia demophon*. a) Karyotype of a spermatogonium, which displays 10-11 sub-metacentic and 5-6 acrocentric pairs. b) Spontaneous pseudo C-banding of a spermatocyte II with compacted proximal and fuzzy distal regions (2 chromosomes are missing). As in the karyotype, acrocentric chromosomes (A) are a minority.

### *Melinaea menophilus* n. ssp

An intra-specific polymorphism is present in karyotypes of the males studied: 38,ZZ and 40,ZZ. Observed mitotic cells, which have long chromosomes with coalescent chromatids, were probably in pro-metaphase and chromosome morphology was hardly visible. The polymorphism is confirmed at metaphase I of meiosis by the variation of the number of bivalents (16 or 17) and the presence of trivalents, which indicates the heterozygote status for two chromosome rearrangements (Fig. 7a). Metaphases II display typical sub-metacentric and acrocentric chromosomes (Fig. 7b). Such morphologies are hardly incompatible with an inverted meiosis.

**Fig. 7.**
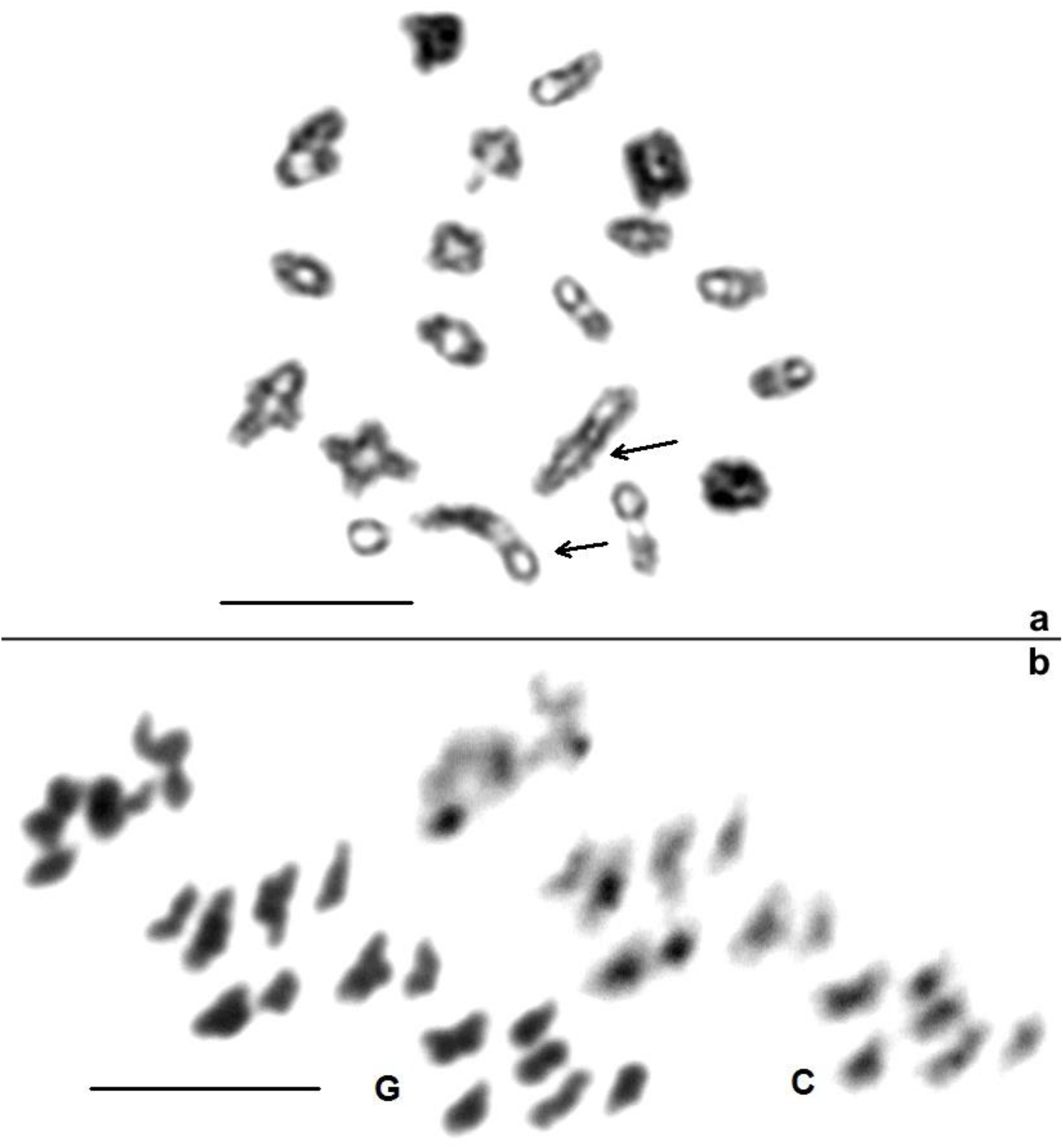
*Melinea menophilus*. a) Metaphase I displaying 16 bivalents and 2 trivalents (arrows). b) Sequentially Giemsa stained and C-banded metaphase II in which many chromosomes look sub-metacentric.

### Ithomia salapia aquinia

The male mitotic karyotype is composed of 68-70 chromosomes. After C-banding, many chromosomes exhibit one band, most often in a terminal region (Fig. 8a). Diakineses display 32 bivalents and 2 trivalents. As usual in our hands with insects, C-banding is more intense in meiotic than in mitotic cells and thus, many more chromosomes look C-banded at diakinesis (Fig. 8b). Individual chromosomes from each bivalent display 0, 1 or 2 C-bands. Differences between homologs (arrow heads) demonstrate the existence of a polymorphism of C-banded heterochromatin repartition. Most bivalents exhibit 1 chiasma in median position and do not differ from acrocentric bivalents of canonical meiosis. A few other bivalents form a distal chiasma, and each homolog exhibits a C-band in median position. On the whole, the location of C-bands at chromosome ends, rather characterizes telomeric than centromeric heterochromatin.

**Fig. 8.**
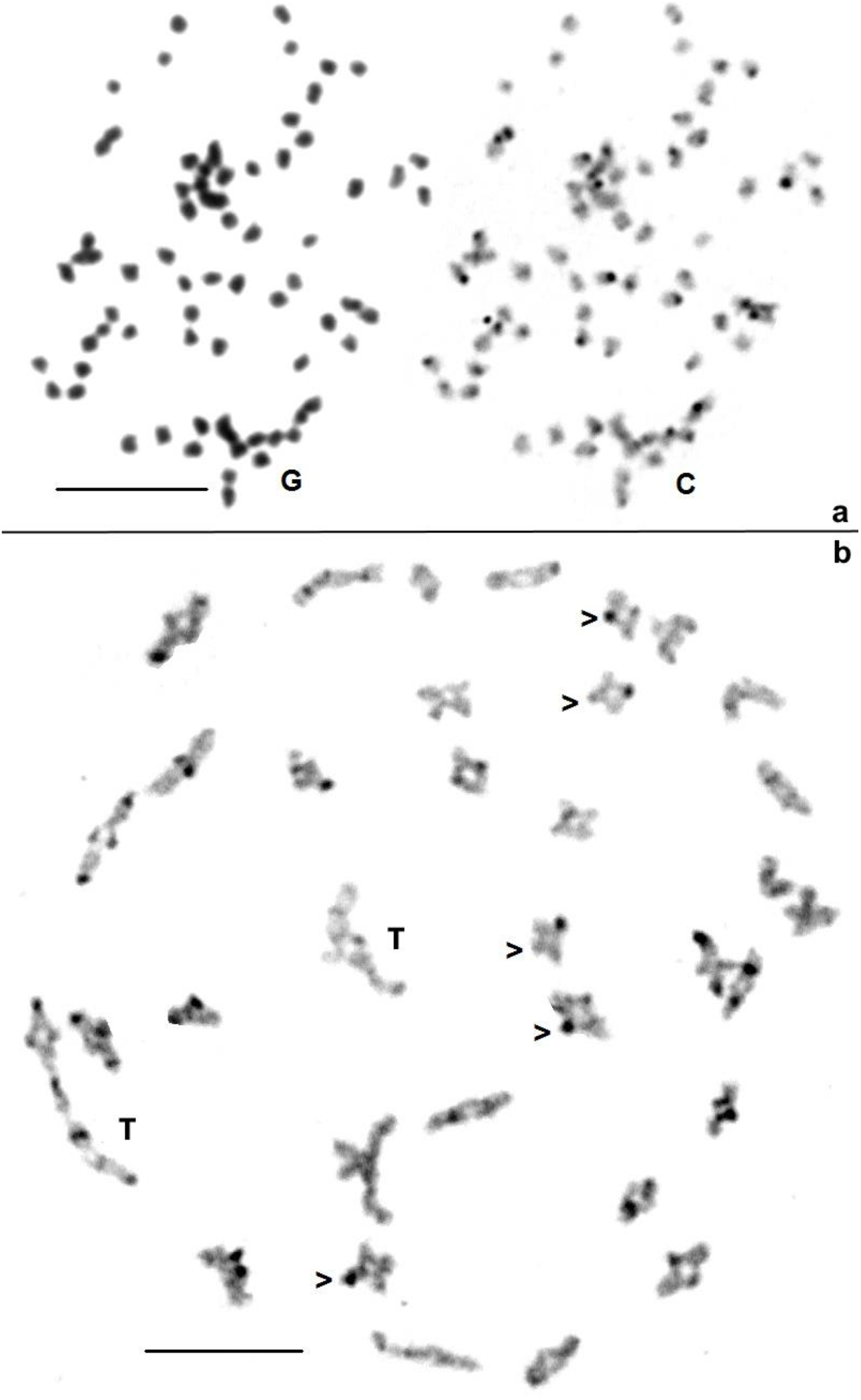
*Ithomia salapia aquinia*. a) Giemsa stained (left) and C-banded (right) spermatogonial metaphase. b) C-banded metaphase I exhibiting 33 bivalents and 2 trivalents (T). C-bands are more intense than in the spermatogonium and located to up to 4 telomeric regions per bivalent. Their frequent asymmetry (arrow heads) demonstrates a strong polymorphism. They are located at intercalary position in a few bivalents, possibly formed by 2 metacentric chromosomes.

## Discussion

### Mitotic chromosomes exhibit atypical centromeric regions

In a recent study of genus *Melinaea* (Nymphalidae) in which presumably holocentric chromosomes were involved in a complex evolution by multiple fissions-fusions, we were surprized to observe that each chromosome could be attached to a single and not to multiple spindle fibres, as would be expected for holochromosomes at anaphase I of meiosis [McClure et al, 2018]. However, the lack of clear primary constriction on their mitotic chromosomes prevented us to challenge their holokinetic nature. By using improved techniques, the classification of mitotic chromosomes based on their “acrocentric” or “sub-metacentric” morphology is however possible in species with low chromosome numbers. Indeed, this suggests that chromosomes are not holocentric, but compared to classical mono-centric chromosomes their morphology is particular: they do not exhibit the Giemsa negative primary constrictions that mark centromeric regions. Usually, these primary constrictions harbour, in addition to the proper centromere, large amounts of highly repeated (satellite) DNA resistant to denaturation and nested into so-called constitutive heterochromatin generally stained after C-banding. The putative centromere regions of butterflies shown above are marked only by a well localized, but loose coalescence of the chromatids. The poorness in highly repeated sequences (satellite DNA) in the genome of butterflies may explain the frequent paucity of heterochromatin, C-banding and primary constrictions, but it does not demonstrate the lack of functional centromeres, which may be nested in other DNA sequences, more difficult to identify. At metaphase/anaphase transition, the use of a high-resolution confocal microscope on DAPI stained chromosome disclose discreet chromatin, which links the two sister chromatids at the place of the putative centromere (Fig. 4). Yet, butterfly chromosomes lack the centromeric protein CenH3 (Cenp-A of mammals), considered to be present in all centromeres of monocentric chromosomes [Drinnenberg et al., 2014]. Nevertheless, our observations suggest that butterfly centromeres conserved some functionality despite the loss of CenH3.

The difficulty for distinguishing butterfly chromosome morphology is materialised by comparing the (2N = 8) karyotype of *D. melanogaster* to the (2N = 50) karyotype of *P. rapae*, which has a slightly higher DNA content (Table 1). The DNA of an average chromosome of *P. rapae* is composed of about 9.8 Mb, whereas that of chromosome 2 of D. melanogaster is composed of 48.8 Mbp, hence a ratio of 1/5th. Likewise, the average DNA-content of a medium sized butterfly chromosome represents about 8% of human chromosome 1. According to ISCN [1985], this chromosome exhibits about 27 bands at metaphase (M), 40 bands at pro-metaphase (PM) and 50 bands at prophase (P). An average butterfly chromosome is scarcely larger than 2 (M), 3 (PM) or 4 (P) bands and is therefore very small, compared to a mammalian chromosome. This may explain why, in the literature, butterfly chromosomes usually look like dots at metaphase and thin filaments with coalescent chromatids at prophase. As in any prophase, their chromatids are hardy discriminated. Furthermore, their centromeres, devoid of heterochromatin, are hardly detectable. Numerical variations of chromosome may be also used to challenge their holocentricity, by comparing closely related species with very different chromosome numbers, as the 2 Pierids studied here. Proportionally, more sub-metacentric chromosomes are expected in karyotypes with a smaller number of chromosomes than values close to the presumed ancestral number (i.e. 60-62). Indeed, sub-metacentric chromosomes are observed in *P*.*brassicae* (2N = 30) and *A. demophon* (2N = 32), but hardly in *P. rapae* (2N = 50) or A. eurypyle (2n = 62) karyotypes, in which acrocentric chromosomes largely predominate. In butterfly species with such high chromosome numbers, many acrocentric chromosomes are expected and their heterochromatin poor centromeres come near or at contact with telomeric heterochromatin [Schoeftner and Blasco, 2009, Chawla and Azzalin, 2008], whose instability [Murname, 2012] may be responsible of their multiple rearrangements, notably fusions. Another factor possibly involved in butterfly chromosome evolution is the very frequent distal position of their 18S rDNA genes on one or multiple chromosomes [Provaznikova et al., 2021]. In both beetles and Primates, we showed that the terminal position of rDNA genes, which behave like fragile sites, is a major source of transmissible chromosome rearrangements [Dutrillaux et al., 2016, Gerbault-Seureau et al., 2017].

### Meiosis is not inverted

The large intra- and inter-specific numerical variations of butterfly chromosomes usually represent indirect arguments in favour of their holocentricity, supposed to facilitate the correct segregation of asymmetrical chromosomes via an inverted meiosis [Mola and Papeschi, 2006, Lukhtanov et al., 2018]. One particularity of holocentricity is the attachment of chromosome bivalents to multiple spindle fibres at metaphase I/anaphase I of meiosis. A second particularity is that the first division is equational (separation of the two chromatids of each homolog), instead of reductional (separation of the homologs with non-cleaved centromeres). The cytological consequences should be visible at each stage of the meiotic process from the end of diplotene to metaphase II (see Fig. 9 in [Lukhtanov et al., 2018]). We have no argument to contest the existence of such meiosis in some species, but it certainly does not occur in the butterflies studied here for the following reasons:

- diakineses/metaphases I (Fig. 7a, 8b) display all the features of a canonical meiosis;
- metaphase I-anaphase I transition, as shown for *P. rapae* (Fig. 5b) perfectly fits with the presence of acrocentric chromosomes, as also exhibited by mitotic cells: the numerous acrocentrics have a well-marked centromere and clearly separated chromatids and homologs form bivalents in end-to end (chiasmatic or achiasmatic?) association. This configuration is typical of a canonical meiosis;
- metaphases II display monocentric chromosomes, typical for this phase, with strongly apart and fuzzy chromatids linked by a compact centromeric region, often darker than chromatids (Fig. 3c, 6b and 7b). This mimics a C-banding, a feature frequently observed in metaphases II of beetles (data not shown);
- and indeed, in a given species, similar proportions of acrocentric and sub-metacentric chromosomes can be recurrently observed in different mitotic karyotypes (compare Fig. 3a, Fig. 3b and complementary data) and at both mitotic metaphase and metaphase II of meiosis (compare Fig. 3a and 3c).

Finally, the example of *M. menophilus*, which belongs to a genus with very high intra- and inter-specific numerical polymorphisms [McClure et al., 2018], shows that such polymorphisms do not preclude gametogenesis with a canonical meiosis.

## Conclusion

The very small size of butterfly chromosomes makes the analysis of their morphology difficult. This difficulty is worsened by the lack of primary constrictions, i. e., centromeric heterochromatin, which contains highly repetitive (satellite) DNA sequences and is usually associated with particular staining, compaction and joining of sister chromatids. In butterflies, heterochromatin is scarce, and rather more telomeric than centromeric. In such conditions, centromeres are marked by discrete joining of sister chromatids only, more visible in cohesin less meiotic metaphases II than in mitotic pro-metaphases. We show that karyotypes with about 60 chromosomes, a number considered to be close to that of butterfly ancestor, seem to be essentially composed of acrocentrics. Consequently, centromeres of butterfly chromosomes are often located in or at close contact with telomeric heterochromatin, whose composition differs from that of centromeric heterochromatin [DeBaryshe and Pardue, 2011] and whose instability may be responsible of their numerous rearrangements. In other words, the large numerical variations of chromosome observed in butterflies may be better explained by their telomere/centromere instability than by the facilitated transmission of rearranged chromosomes through inverted meiosis, whose occurrence is challenged here.

## Acknowledgments

We thank the Centre de Microscopie de fluorescence et d’IMagerie numérique (CeMIM) from the analytical platform (PAM) at the National Museum of Natural History (MNHN) in Paris, France, for providing access to the confocal laser microscope. We thank SERFOR for providing research permits in Peru, and local assistants.

## Statement of Ethics

The authors have no ethical conflicts to disclose. Ethical approval is not required for this type of research.

## Conflict of interest statement

Les authors declare that they have no conflict of interest.

## Funding sources

This research was funded by the Agence Nationale de la Recherche (grants SPECREP and CLEARWING) and by a HSFP research grant (RGP0014/2016)

## Author contributions

Bernard and Anne-Marie Dutrillaux conceived the study and performed the technical work. Marianne Elias obtained the funding. Bernard Dutrillaux, Anne-Marie Dutrillaux, Mélanie McClure and Marianne Elias obtained the samples. Marc Gèze performed the confocal study. All authors discussed the project and findings. Bernard Dutrillaux wrote the manuscript with extensive contributions from Bertrand Bed’Hom and Marianne Elias. All authors read and approved the manuscript.

## Data availability statement

All data are included in the manuscript and in supplementary data

## Complementary data

Fig.S1, S2, S3 and S4. Other karyotypes of *P. brassicae* after Giemsa staining (ganglia cells, Fig.S1 and S2), C-banding (Fig.S3, egg) and spermatocyte I at anaphasic segregation (Fig.S4). The almost entirely heterochromatic chromosome (Fig.S3) may be a W. All other C-bands are polymorphic and located at chromosome end

## Notes

### Competing Interest Statement

The authors have declared no competing interest.

