## Supplementary figures for "Improved basic cytogenetics challenges holocentricity of butterfly chromosomes"

a. Institut de Systématique, Evolution, Biodiversité (ISYEB), Muséum National d'Histoire Naturelle, CNRS, Sorbonne Université, EPHE, Université des Antilles, Paris, France
b. Laboratoire Ecologie, Evolution, Interactions des Systèmes Amazoniens (LEEISA), Université de Guyane, CNRS, IFREMER, Cayenne, France
c. Molécules de Communication et Adaptation des Micro-organismes (MCAM), Muséum National d'Histoire Naturelle, CNRS, Paris, France
d. Centre de Microscopie de fluorescence et d'Imagerie numérique (CeMIM), Muséum National d'Histoire Naturelle, Paris, France


**Supplementary figures**

Fig.S1, S2, S3 and S4. Other karyotypes of *P. brassicae* after Giemsa staining (ganglia cells, Fig.S1 and S2), C-banding (Fig.S3, egg) and spermatocyte I at anaphasic segregation (Fig.S4). The almost entirely heterochromatic chromosome (Fig.S3) may be a W. All other C-bands are polymorphic and located at chromosome ends.

Figure S1

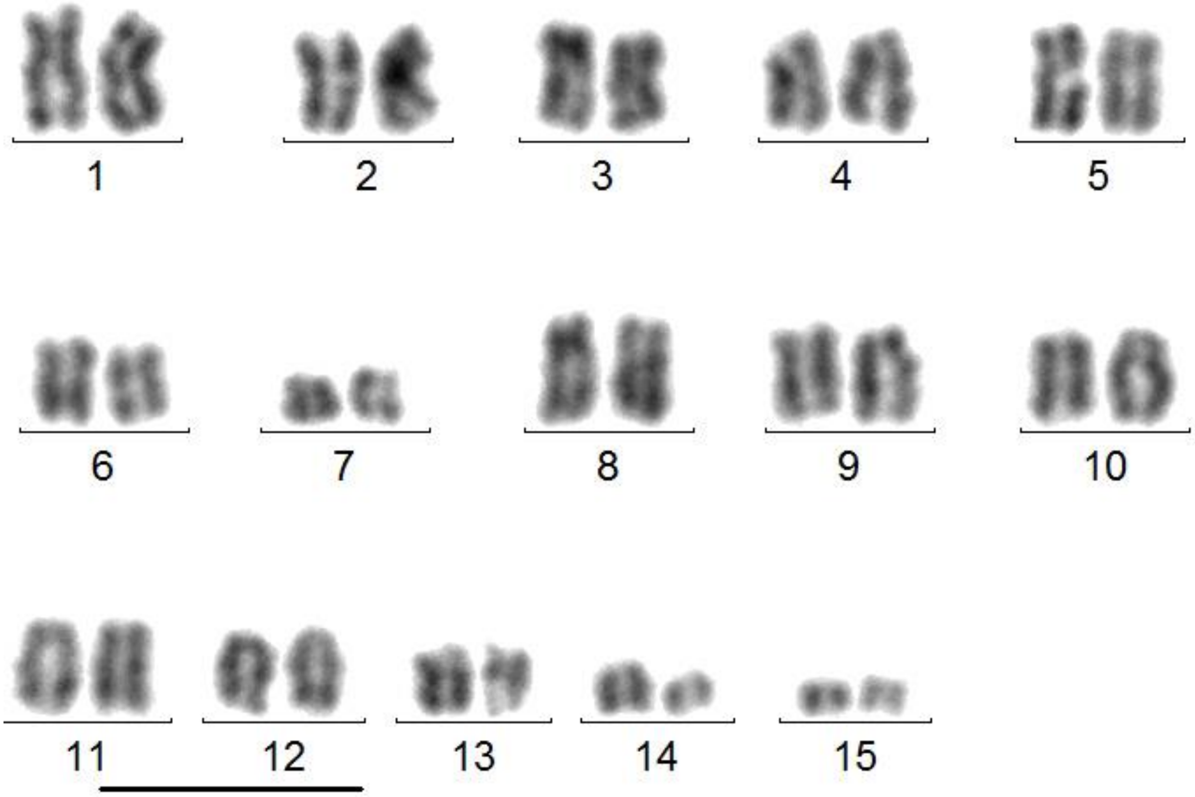


Figure S2

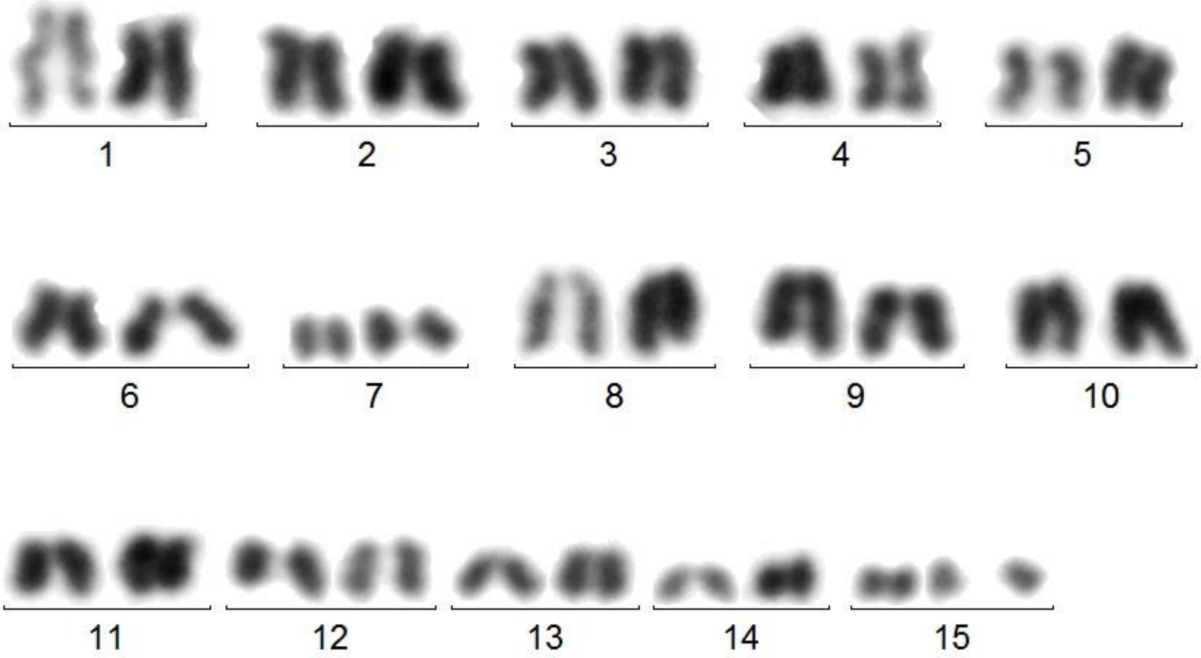

Figure S3

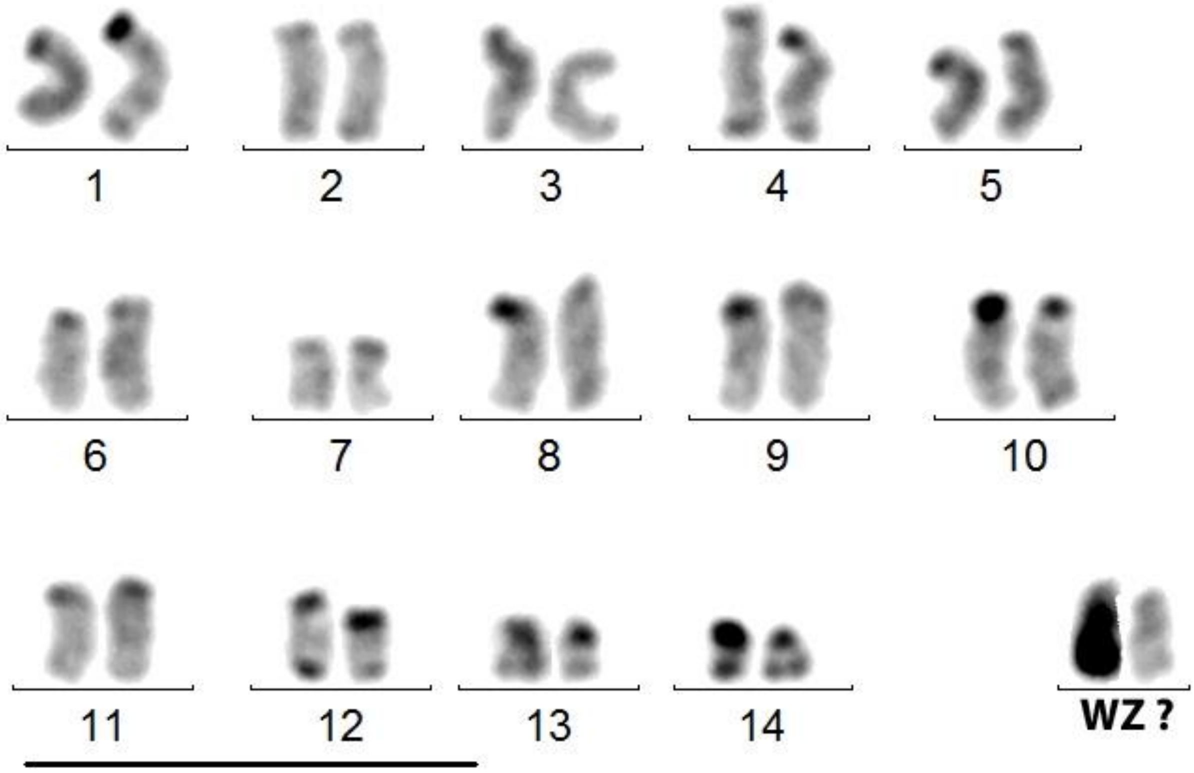


Figure S4

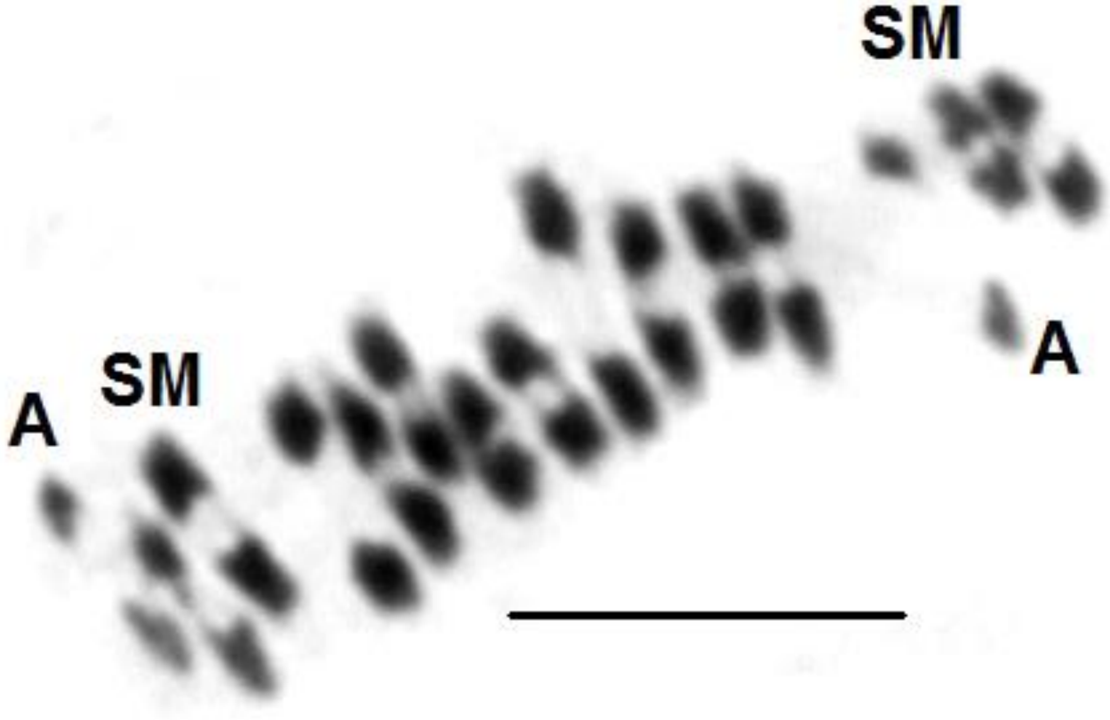
